## Supplementary file for "Experimental evolution of a pathogen confronted with innate immune memory increases variation in virulence"

**Table 1** qPCR primers and their amplification efficiencies

| gene | forward primer | reverse primer | efficiency |
|---|---|---|---|
| *Cry3a* | AACAGATGAAGCAAGTACACAAACG | CTGTTGTTTCTGGAGGCAATTGATC | 105% |
| *Yqey* | AGCTGGTCGTGAAGACCTTG | CGGCATAACAGCAGTCATCA | 94% |
| *rps21* | AAATCTGAAGCGGCAAGAA | AAGATCGGTTTCTAAACTGGTACA | 102% |

**Table 2** Relative expression of *Cry3a* gene in cultures from different *Btt* strains. *Btt*- and *Bt407*- do not carry the *Cry3a* gene. Priming and control lines were evolved from ancestral strain. Mean ΔCt values by subtracting the target gene Ct value from the geometric mean of the Ct values of the housekeeping genes.

| bacterial culture | mean ΔCt | std |
|---|---|---|
| Btt- | -11.19 | 3.69 |
| Bt407- | -10.15 | 0.57 |
| priming | 0.62 | 1.38 |
| control | 1.01 | 1.31 |
| ancestral | 1.77 | 1.31 |

**Table 3** Differences of *Cry3a* gene expression of individual evolved lines compared to the ancestral strain. Shown are the results of linear mixed effects model with block as experimental factor. Significant differences are shown in bold type face.

| line | estimate | s. e. | dF | t-value | p-value |
|---|---|---|---|---|---|
| intercept | 1.77 | 0.52873 | 5.62228 | 3.342 | 0.017 |
| C1 | -0.56 | 0.59026 | 50 | -0.947 | 0.348 |
| **C2** | **-1.23** | 0.59026 | 50 | -2.087 | **0.042** |
| C3 | 0.25 | 0.59026 | 50 | 0.421 | 0.675 |
| C4 | 0.52 | 0.59026 | 50 | 0.878 | 0.384 |
| **C5** | **-2.34** | 0.59026 | 50 | -3.958 | **<0.001** |
| **C6** | **-1.52** | 0.59026 | 50 | -2.573 | **0.013** |
| C7 | 0.31 | 0.59026 | 50 | 0.525 | 0.602 |
| **C8** | **-1.48** | 0.59026 | 50 | -2.514 | **0.015** |
| P1 | -0.04 | 0.59026 | 50 | -0.062 | 0.951 |
| P2 | -0.71 | 0.59026 | 50 | -1.203 | 0.235 |
| P3 | -0.83 | 0.59026 | 50 | -1.412 | 0.164 |
| P4 | -0.74 | 0.59026 | 50 | -1.26 | 0.213 |
| **P5** | **-2.44** | 0.59026 | 50 | -4.136 | **<0.001** |
| **P6** | **-3.52** | 0.59026 | 50 | -5.964 | **<0.001** |
| **P7** | **-1.52** | 0.59026 | 50 | -2.575 | **0.013** |
| P8 | 0.65 | 0.59026 | 50 | 1.101 | 0.276 |

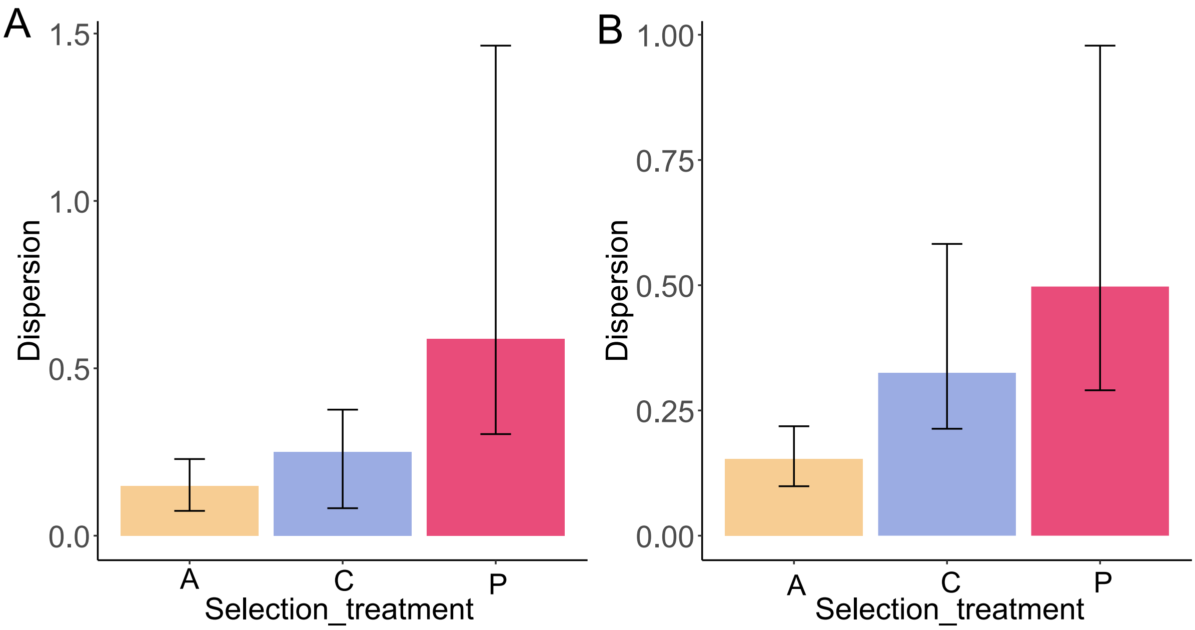


**Figure S1.** Dispersion host mortality in **A** control and **B** primed larval environments. Using a Beta distribution in a Bayesian framework we allowed the selection treatments to influence the precision parameter (φ). By estimating the degree of dispersion (1/ φ) we were able to compare if the variation in virulence was higher in primed evolved bacteria. We found that the variation in virulence (i.e., dispersion host mortality) was greater in primed-evolved *Btt* than in control-evolved *Btt* with a certainty of 99.06%. This probability slightly decreased to 90.14% in primed host environment

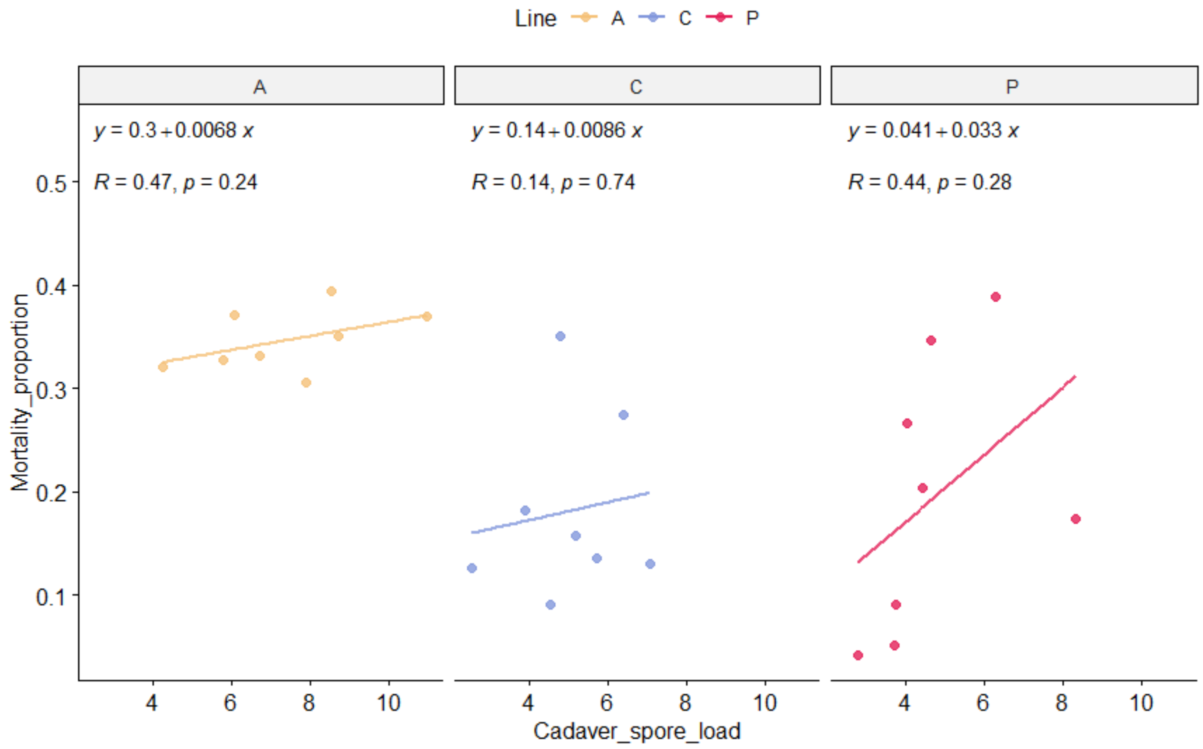


**Figure S2.** Correlation between spore load and virulence in ancestral line (A), control line(C) and primed line (P) in primed hosts

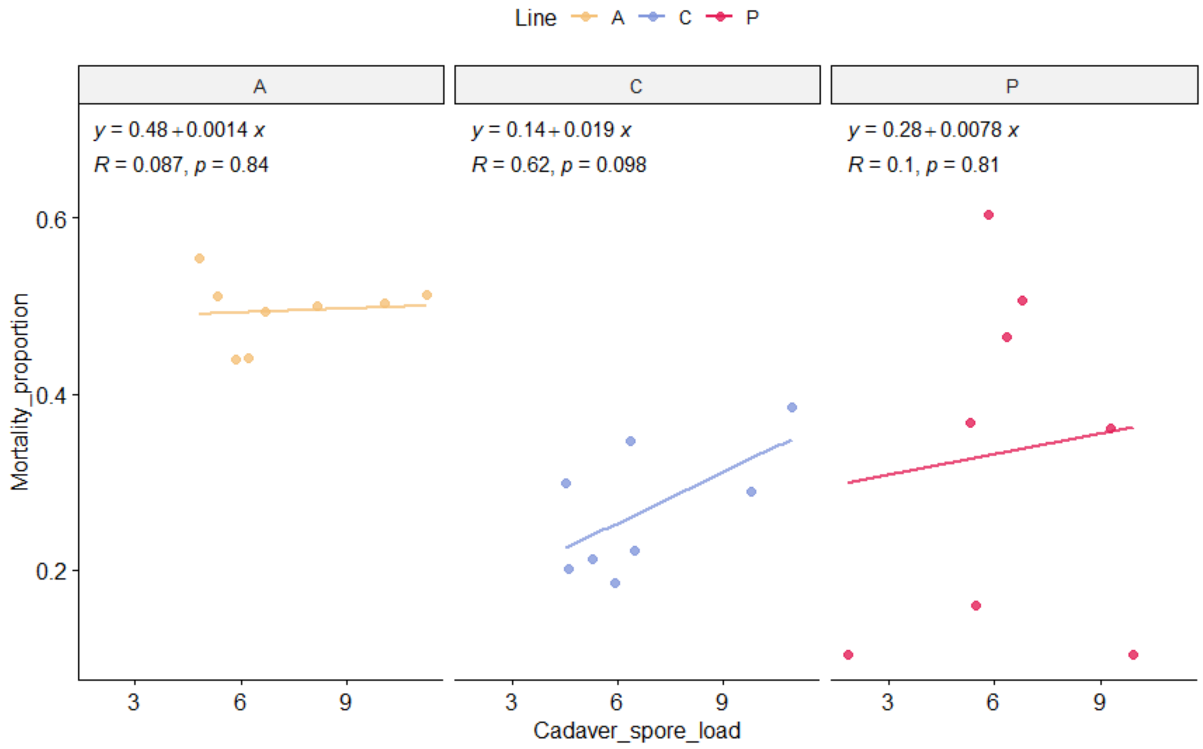


**Figure S3.** Correlation between spore load and virulence in ancestral line (A), control line(C) and primed line (P) in primed beetle larvae in control host

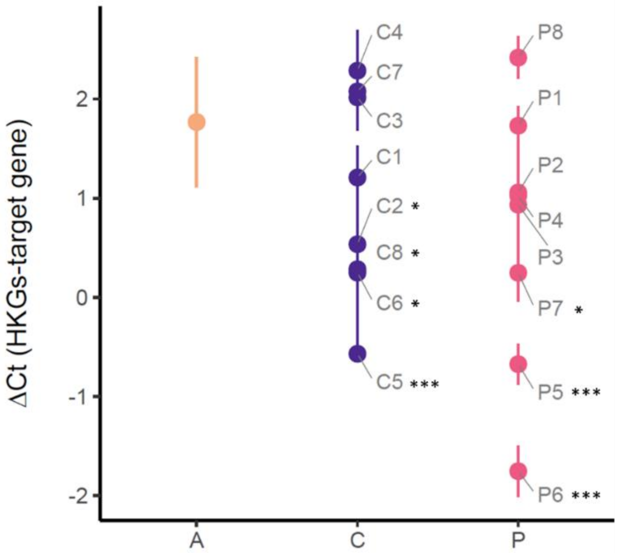


**Figure S4.** Relative expression of *Cry3a* gene for evolved lines. ΔCt values were calculated by subtracting the Ct value of the target gene, *Cry3A* from the geometric mean of the Ct values of two housekeeping genes (*Yqey* and *rps21* ). Per evolved line and ancestral strain, the mean and standard error for four replicates are given. Asterisks indicate significant differences to the ancestral strain (*= p-value <0.05, ***= p-value <0.0001)

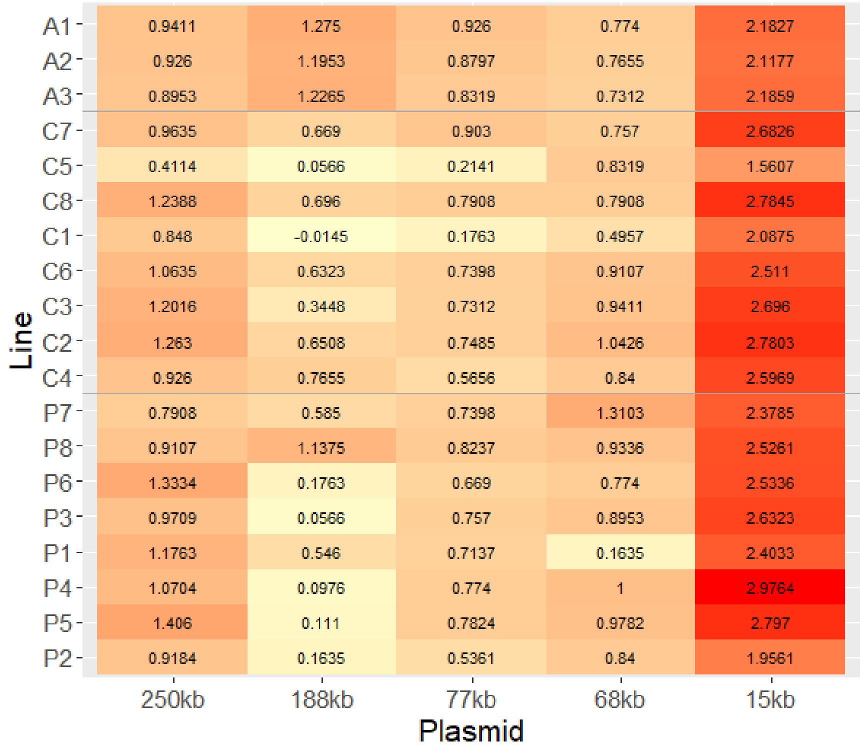


**Figure S5:** Heatmap for the Log2 transformed values of plasmid coverage divided by chromosome coverage for each replicate line. A = ancestral, C = control evolved, P = primed evolved. The evolved lines are ordered by evolution treatment and by virulence.

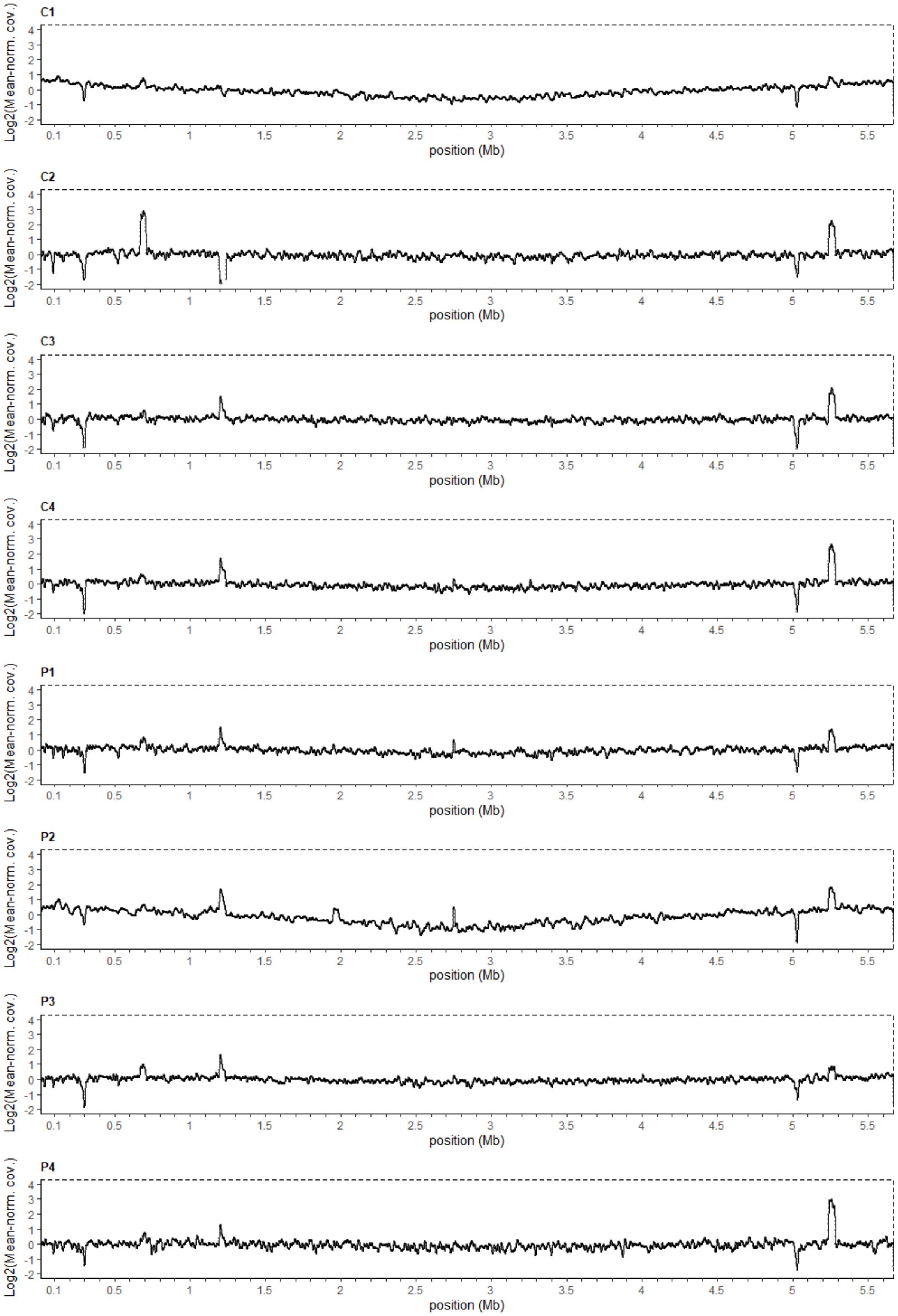


**Figure S6:** Exemplary log2 transformed mean-normalized coverage plots of reads across the chromosome for control (C1 – 4) and priming (P1 – 4) evolved lines.

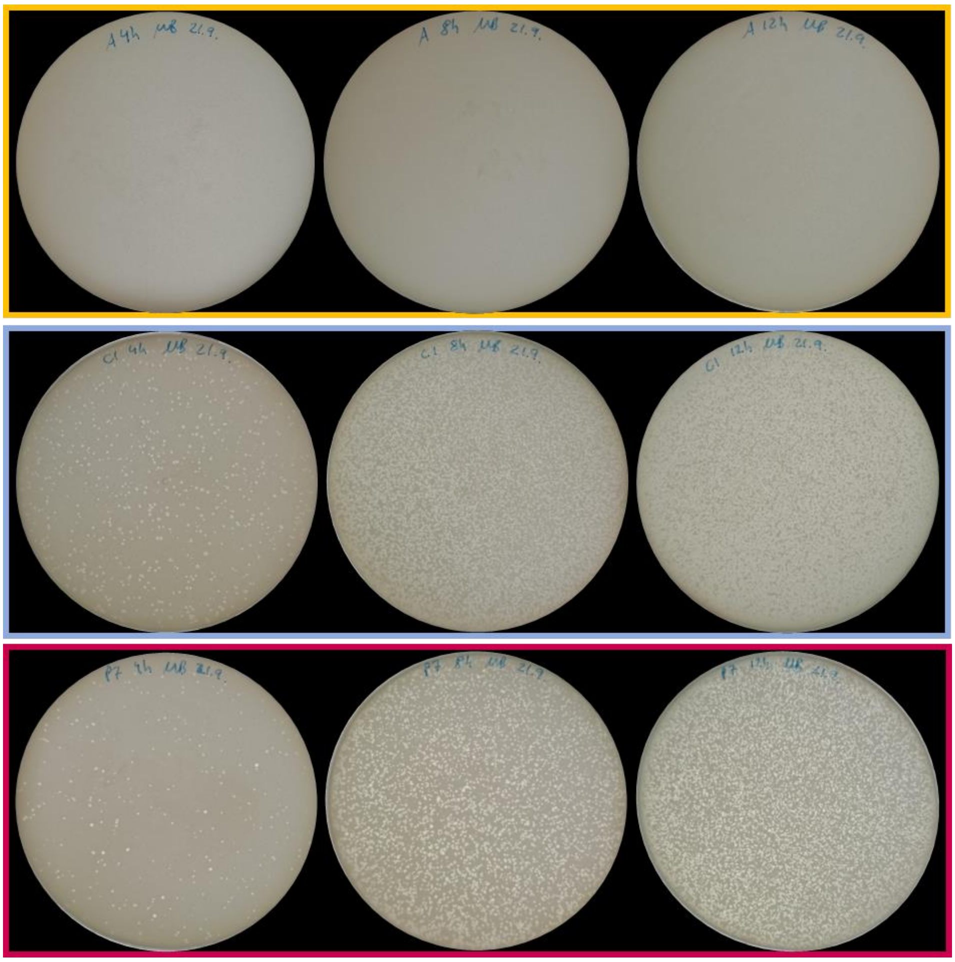

**Figure S7:** Double layer agar assays of an ancestral line A and evolved lines C1 (control evolved) and P7 (priming evolved). Culture samples were taken after 4 hours, 8 hours and 12 hours of growth in LB medium.

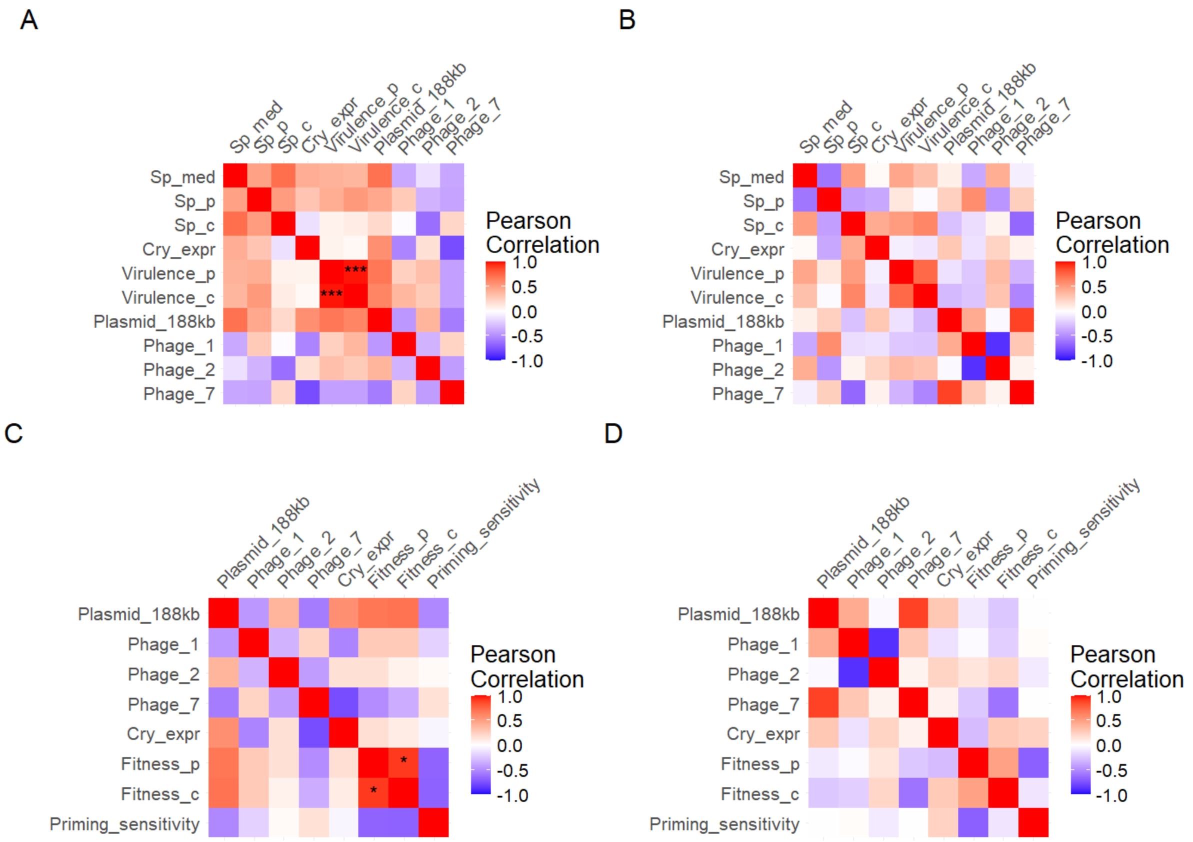


**Figure S8:** Pearson correlation matrix for the primed evolved pathogen (A, C) and the control evolved pathogen (B, D). Correlated traits include all phenotypic traits with active mobile elements (A, B) or Fitness and Priming sensitivity values with active mobile elements (C, D). Asterix indicates which correlations were statistically significant (FDR adjusted p value < 0.05). None of the correlations were statistically different between the primed and control evolved pathogen (BH adjusted p value < 0.05).
